# Rotavirus pre-symptomatically downregulates ileum-innervating sympathetic nerves concomitant with increased intestinal transit and altered brain activity

**DOI:** 10.1101/2021.08.06.455368

**Authors:** Arash Hellysaz, Lennart Svensson, Marie Hagbom

## Abstract

While diarrhea, the hallmark symptom of rotavirus infection, has been considered to occur only due to intrinsic intestinal effects, we show evidence for central control underlying the symptomology. With large-scale 3D volumetric tissue imaging a mouse model, we show that rotavirus infection disrupts the autonomic balance by downregulating the noradrenergic sympathetic nervous system in ileum, concomitant with increased intestinal transit. A most interesting observation was that nervous response from CNS occurs pre-symptomatically, an observation that bring new understanding to how virus give raise to clinical symptoms. In the CNS of infected animals, we found increased pS6 immunoreactivity in the area postrema and decreased phosphorylated STAT5-immunoreactive neurons in the bed nucleus of the stria terminalis, which are associated with autonomic control including stress response. Our observations bring new and important knowledge of how rotavirus virus infection induce gut-nerve-brain crosstalk giving raise to sickness symptoms.

## Introduction

Rotavirus is the major cause of paediatric gastroenteritis, resulting in acute diarrhoea and vomiting. In 2019, rotavirus was estimated to have caused more than 150,000 dehydration-associated child deaths and the hospitalization of millions of children younger than 5 years old (Debellut et al., 2021). The disease mechanisms behind rotavirus-induced diarrhoea and vomiting are still not fully understood and no symptomatic treatment are available. While it is well established that diarrhoea and vomiting are the hallmarks of rotavirus infections, the extent of infection and the involvement of the central nervous system (CNS) in the illness have remained elusive.

Rotavirus non-structural protein 4 (NSP4) stimulates the enterochromaffin (EC) cells of the small intestine to release serotonin (Bialowas et al., 2016; Hagbom et al., 2011), which is sensed by neurons and leads to direct and indirect activation of both the enteric and central nervous systems (ENS and CNS, respectively). Consequently, it has been suggested that vomiting is elicited by gut–brain cross-talks involving the ascending and descending vagal pathways relayed through the vomiting centre in the brain (Crawford et al., 2017; Hagbom et al., 2011). Moreover, illness is not only associated with diarrhoea and vomiting, but also triggers other symptoms such as nausea, fever, anorexia and sickness symptoms, revealing a complex mechanism of disease and further indicating the participation of the CNS.

While intrinsic factors of rotavirus-induced diarrhoea have been investigated (Istrate, Hagbom, Vikström, Magnusson, & Svensson, 2014; S. Kordasti, Sjövall, Lundgren, & Svensson, 2004; Ove Lundgren et al., 2000) the role of CNS in rotavirus illness symptoms remain uncharted. Although the ENS drives intestinal motility independently (Wood, Alpers, & Andrews, 1999), it is *de facto* modulated centrally by the autonomic and endocrine nervous systems (Browning & Travagli, 2014). The inhibitory and excitatory effects of the autonomic nervous system on the small intestine through the sympathetic and the parasympathetic systems are well established (O. Lundgren, 2000; Sharkey & Pittman, 1996; Wood et al., 1999). Normal conditions are defined by the proper balance between these two opposing systems, and balance disruption by either up- or downregulation in either system can disrupt proper motility control and lead to either diarrhoea or constipation.

Recent developments in tissue clearing techniques, such as iDISCO (immunolabeling-enabled three-dimensional imaging of solvent-cleared organs) (Renier et al., 2014), together with volumetric imaging of large samples with light-sheet microscopy (Fadero et al., 2018) and computer-aided analysis of big data, have enabled 3D organ-wide investigation. Here, we used these techniques to study the extent of organ-wide rotavirus infection. Furthermore, we used the same techniques to investigate the effect of rotavirus infection on the sympathetic innervation and activity of the infected small intestine in ways previously not possible. We demonstrate that rotavirus infection of the small intestine pre-symptomatically disrupts the autonomic balance by downregulating the noradrenergic sympathetic nervous system in ileum, concomitant with increased intestinal transit.

## Methods

### Animals

Five to seven–day-old neonatal mice of both sexes and 8–10-week-old female adult BALB/c mice were used. All animal experiments had been approved by the local ethical committee in Linköping, Sweden (approval no.: N141/15 and 55-15).

### Rotavirus infection

The mice were orally infected with 100 diarrhoea doses (100_DD_) of EDIM rotavirus in 10 µL 0.9% saline as described previously (Hagbom et al., 2011; Istrate et al., 2014). Non-infected control mice were mock-infected with 10 µL 0.9% saline. The groups were kept in separate litters, and whole litters were infected simultaneously and housed with their mother during the entire experimental period.

### Tissue preparation

For iDISCO, segments of the small intestine were placed in 4% formaldehyde at room temperature for 24 h, and then transferred to phosphate-buffered saline (PBS) and stored at 4°C until tissue clearing was started.

For immunofluorescence, infant mice were sacrificed, and the brains were resected and fixed for 48 h in 4% formaldehyde solution (Histolab, Sweden). Adult animals were perfused, and their brains resected and fixed for 2 h in 4% formaldehyde. Subsequently, the brains were transferred to 15% sucrose in PBS for 7 days at 4°C, then rapidly frozen and stored at -80°C until sectioning was performed. The brains were cut into 14-µm thick sections on a cryostat (Microm; Walldorf, Germany), mounted on chrome alum gelatin-coated slides and stored at -20°C for subsequent immunofluorescence processing.

### Immunofluorescence

The slides were thawed to room temperature, incubated in PBS, and processed for conventional indirect immunofluorescence or tyramide signal amplification (TSA; Perkin Elmer, Waltham, MA, USA) protocols as described previously (Foo, Hellysaz, & Broberger, 2014). All reactions were performed at room temperature unless otherwise stated. Primary antisera cocktails were prepared in staining buffer containing 0.03% Triton X-100 in 0.01 M PBS with 1% bovine serum albumin at least 24 h before use.

For conventional immunofluorescence, the sections were incubated in primary antisera at 4°C for 16 h, rinsed in PBS for 30 min, incubated for 1 h in secondary antisera cocktail, diluted in staining buffer, incubated in 4’,6-diamidino-2-phenylindole (DAPI, 1:10,000 in PBS), rinsed in PBS for 30 min, and mounted with 2.5% 1,4-diazabicyclo[2.2.2]octane (DABCO, Sigma, St. Louis, MO, USA) anti-fade agent in glycerol.

For TSA, antigen retrieval was initially performed with Tris-HCl (pH 8.0) at 95°C for 5 min. The sections were subsequently washed in Tris-sodium chloride-Tween buffer (TNT; 0.1 M Tris, 0.15 M NaCl, 0.05% Tween 20), incubated in primary antisera at 4°C for 42 h, washed in TNT, pre-incubated with blocking buffer (TNB) supplied in the TSA Plus kit (Perkin Elmer) for 30 min, incubated for 2 h in secondary antisera cocktail diluted in TNB, rinsed with TNT buffer, and incubated for 10 min with tyramide-conjugated fluorescein (1:500 in amplification diluent as supplied with the TSA Plus kit). The sections were then stained for DAPI and mounted as described above.

### iDISCO

Approximately 5-mm long intestinal tissue samples from the duodenum and ileum were processed for iDISCO (Renier et al., 2014) according to the May 2016 updated protocol (available at http://www.idisco.info/), with some modifications. Briefly, the samples were dehydrated with gradient methanol, bleached in chilled fresh 5% H_2_O_2_ in methanol overnight at 4°C, and rehydrated and washed in PBS with 10 mg/L heparin and 0.2% Tween 20. Subsequently, the samples were permeabilized for 1 day, blocked for 1 day, incubated in primary antibody at 42°C for 7 days, washed, incubated in secondary antisera at 42°C for 7 days, and washed.

Following methanol gradient dehydration, the samples were incubated for 3 h in 66% dichloromethane in methanol, 2 × 15 min in 100% dichloromethane, and transferred to dibenzyl ether. All samples used for final analysis were processed in parallel and treated with the same buffers and solutions.

### FITC-dextran transit

At 16 h p.i., the animals were orally administered 10 µL freshly prepared 4-kDa FITC-dextran (FD-4s, Sigma) dissolved in Milli Q water, at a dose of 0.25 mg/animal. After 15 min, the animals were sacrificed and the entire intestine, from the stomach to the rectum, was removed and visualized with ultraviolet light in a ChemiDoc XRS system (Bio-Rad, Sweden). The front part of the main accumulating FITC-dextran was defined from the photo, and the software program Adobe Illustrator was used to exactly measure the intestinal length and migration of the FITC-dextran probe. Intestinal transit was calculated on how far the FITC-dextran probe has passed as a percentage of the entire length of the intestine, from the pylorus to the rectum (Istrate et al., 2014).

### Antisera

All antisera used in the different protocols are presented in Table 1. For detection, Alexa Fluor-conjugated secondary antisera (Life Technologies, Carlsbad, California, United States) for conventional detection, and horseradish peroxidase-conjugated secondary antisera (Dako, Glostrup, Denmark) for TSA were used. All secondary antisera were diluted to 1:500 for IHC and TSA and 1:250 for iDISCO.

**Table 1.**
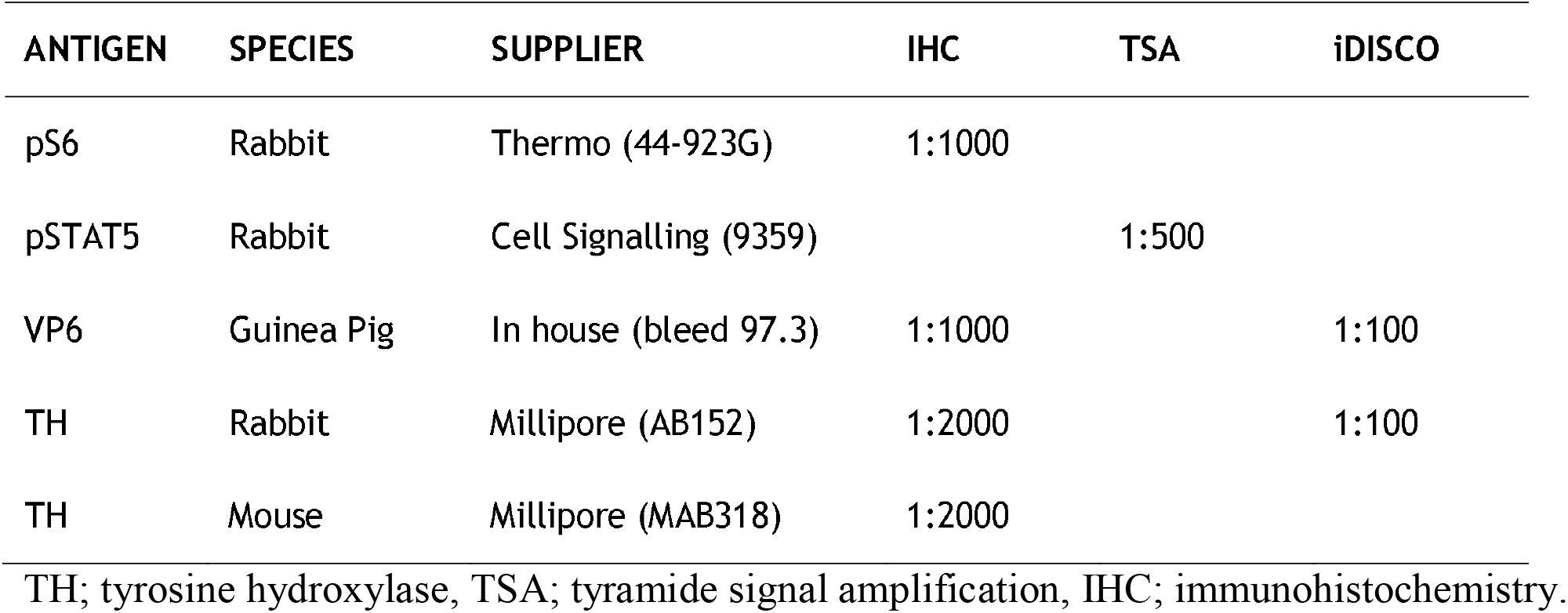
Primary antisera used in the study.

### Microscopy

Wide-field image montages were automatically generated in Neurolucida computer software (MBF Bioscience, Williston, VT, USA) by taking consecutive pictures with an automated stage controller mounted on a Zeiss Axio Imager M1 (Carl Zeiss, Oberkochen, Germany). Confocal micrographs were captured using a Zeiss LSM 800 Airyscan microscope with Zen Blue computer software. Light-sheet micrographs were acquired with a UltraMicroscope II (LaVision Biotec, Bielefeld, Germany) setup using ImSpector computer software. All intestinal tissues were randomized and sampled consecutively with the same acquisition settings. Post-acquisition brightness/contrast adjustments were performed uniformly on all light-sheet micrographs.

### Micrograph analysis

The fluorescence micrographs were post-processed for rotation and brightness/contrast in Photoshop (Adobe, San Jose, California, United States) and analyzed in QuPath computer software (Bankhead et al., 2017).

We performed 3D confocal and light sheet analyses in Imaris. To maintain uniform tissues and measurements between animals, intestinal tissue integrity was visually confirmed in 3D, and damaged segments lacking an intact myenteric plexus (Figure S4) were excluded from analysis. Furthermore, the reconstructed 3D models were trimmed *in silica*, and only fragments with fully intact submucosa and myenteric plexuses were used. Therefore, mucosal immunofluorescence from, for example, enteric dopaminergic cells (Figure 2g) and intense fluorescence from incoming axon bundles (compare Figure 1j, k) were not included in the analysis and did not falsely skew the results.

Two different approaches were used to assess the level of infection in the small intestine. First, the number of infected cells per volume was estimated from the total number of infected surfaces and the total volume of the analyzed tissue. For a more accurate estimation, we set the infected surface creation pipeline to consider cell diameter and split touching objects (see parameters and threshold settings in Figure S5). In the second approach, the tissue infection ratio was estimated based on the total volume occupied by rotavirus relative to the total tissue volume. This approach for estimating the level of infection is independent of cell size and is therefore prone to methodological errors introduced by the splitting algorithm, from which the first approach might suffer from. Both non-infected and infected sample were analyzed with the same analysis pipeline.

### Statistical analysis

Statistical analysis was performed with Prism (GraphPad, San Diego, California, United States) computer software. Statistical significance was set at p < 0.05 and was determined using the statistical tests described in the figures (*p < 0.05; **p < 0.01; ***p < 0.001; ns, not significant). The statistics are reported as the mean ± standard error of the mean (SEM); n corresponds to the number of animals unless indicated otherwise.

### Data availability

Data is available from the corresponding author upon request.

## Results

### Rotavirus infection is widespread throughout the entire length of the small intestine at 16 h post infection

Light-sheet micrograph stacks of the duodenum (Figure 1a, b, Supplementary Video 1) with 3D reconstruction (Figure 1c), and the ileum (Figure 1d, Supplementary Video 2-3) with 3D reconstruction (Figure 1e, Supplementary Video 4), immunostained for rotavirus structural viral protein 6 (VP6), indicated uniform and widespread infection throughout the entire length of the small intestine. VP6 immunoreactivity was not observed in non-infected animals (Figure 1f). Notably, the presence of VP6 was restricted to the mucosa, and no immunoreactivity was observed in the intestinal wall (Figure 1g).

**Figure 1.**
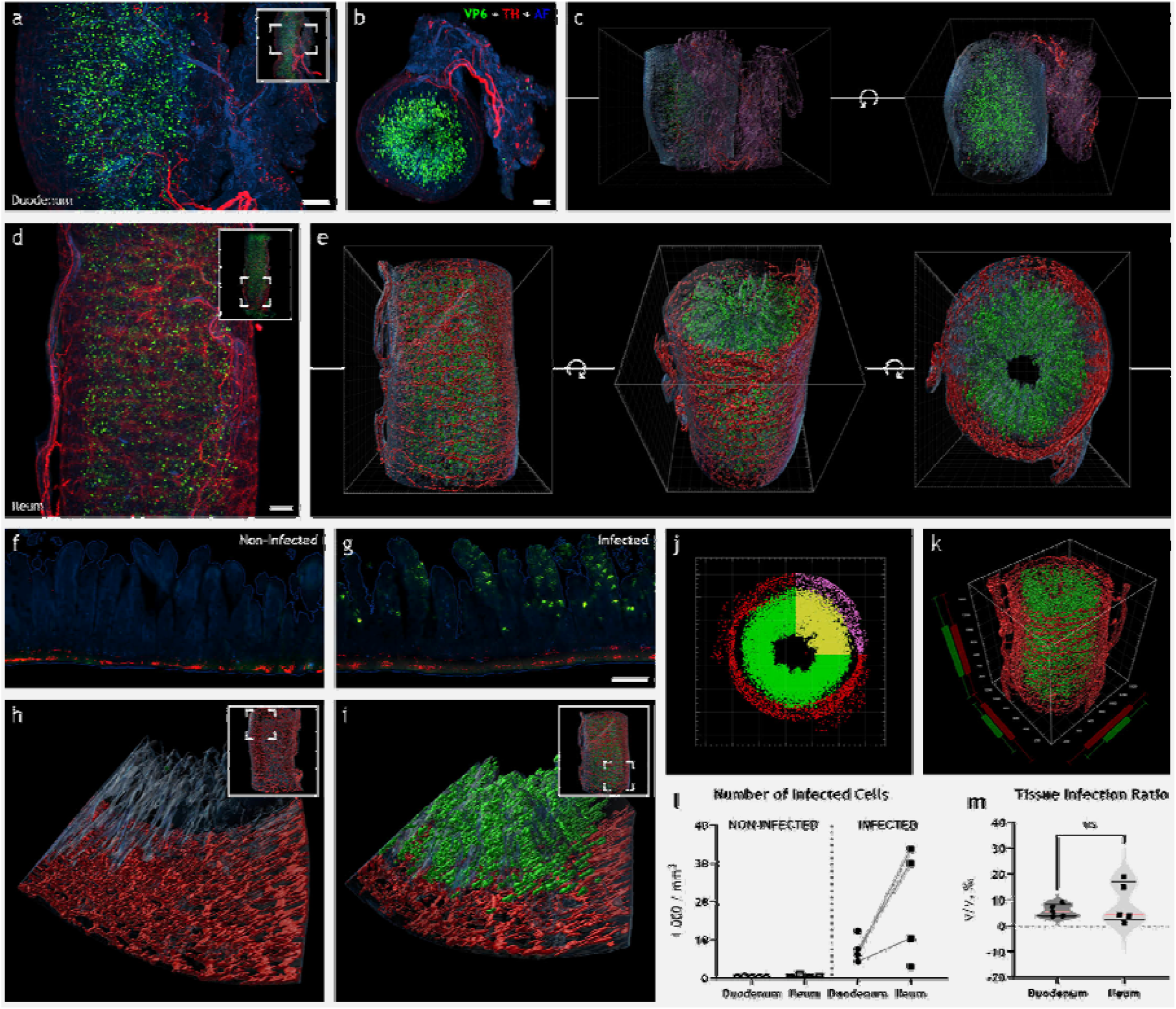
Rotavirus infection is widespr ead throughout the entire length of the small intestine at 16 hours post infection. Maximum intensity projection of light-sheet micrograph stacks (**a, b, d**) from rotavirus-infected mouse duodenum (**a, b**) and ileum (**d**) stained for rotavirus VP6 (green) and TH (red), the rate-limiting enzyme in catecholamine biosynthesis. Tissue was visualized with autofluorescence (AF; blue). Insets show low-power micrographs denoting enlarged regions in the panel with a box. 3D surface reconstruction from (**a, b**) and (**d**) is shown in (**c**) and (**e**), respectively. Rotation along the z-axis is denoted with (↺). Note the high degree of rotavirus infection in the duodenum (**a–c**) and ileum (**d, e**). Single optical slice (**f, g**) and surface 3D reconstruction (**h, i**) of infected (**g, i**) and non-infected (**f, h**) ileum. Imaris vantage plots (**j, k**) from infected ileum. Note the regions used for analysis marked with yellow/purple, excluding, for example, incoming axon bundles. Rotavirus infection was quantified by estimating the relative number of infected cells (**l**) or the tissue infection ratio (**m**). Data points from the same animal are connected with a line. The two-tailed paired *t*-test yielded no significant difference in the relative number of infected cells (**l**; p = 0.1026) or tissue infection ratio (**m**; p = 0.1826). Scale bar in (**a, b, d**) = 50 µm; in (**g**) = 100 µm for (**f, g**).

Next, the extent of infection was investigated. To quantify the level of rotavirus infection, light-sheet micrographs were processed in Imaris (Bitplane, Zürich, Switzerland), and 3D surface models based on voxel fluorescence intensity were automatically created (Figure 1c, e, h–k). The tissue was modelled using autofluorescence. The level of infection was assessed with two different approaches, where number of infected cells (Figure 1l) or tissue infection ratio (Figure 1m) for non-infected (n = 5) and infected (n = 5) duodenum and ileum was estimated. Notably, both approaches generated similar results and yielded the same conclusions (compare Figure S1).

The estimated number of infected cells was 8504 ± 1615 in the duodenum and 17,458 ± 6058 in the ileum (Figure 1l). Likewise, the estimated tissue infection ratio was 5.8 ± 1.0‰ in the duodenum and 8.6 ± 3.5‰ in ileum (Figure 1m). Therefore, our data show no statistically significant differences in the level of rotavirus infection between the duodenum and the ileum.

### Rotavirus infection induces downregulation of the noradrenergic sympathetic neurons in ileum

The main clinical outcome of gastrointestinal rotavirus infection is diarrhoea, which is caused by altered intestinal secretion and motility (Crawford et al., 2017). As both secretion and motility can be modulated by the autonomic nervous system (Browning & Travagli, 2014), we determined whether rotavirus infection would affect the sympathetic nervous afferents innervating the small intestine.

Within the intestinal wall, all tyrosine hydroxylase (TH), *i*.*e*., the rate-limiting enzyme of noradrenalin biosynthesis (Levitt, Spector, Sjoerdsma, & Udenfriend, 1965; Nagatsu, Levitt, & Udenfriend, 1964), reside within the axons of the sympathetic neurons, and extrinsic sympathetic denervation of the ileum abolishes all traces of TH (Mann & Bell, 1993). We measured the total TH immunoreactivity in 3–4 mm long pieces of the intestinal wall with volumetric 3D imaging (see Supplementary Video 1-7) to assess the extent of sympathetic modulation of the rotavirus-infected small intestine.

Surprisingly, in non-infected animals (Figure 2a–d), we observed a clear difference in TH immunoreactivity between the duodenum and ileum. This difference was not obvious in the infected animals (Figure 1e–h). The measured fluorescence intensity (Figure 1i) was 2155 ± 89 au/µm^3^ in the duodenum (n = 5) and was significantly higher (4601 ± 483 au/µm^3^) in the ileum (n = 5) of the non-infected animals. In the duodenum of the infected animals (n = 5), the fluorescence intensity was 2319 ± 128 au/µm^3^. Accordingly, no significant differences in TH immunoreactivity could be observed in the duodenal wall of the infected *vs*. non-infected animals.

**Figure 2.**
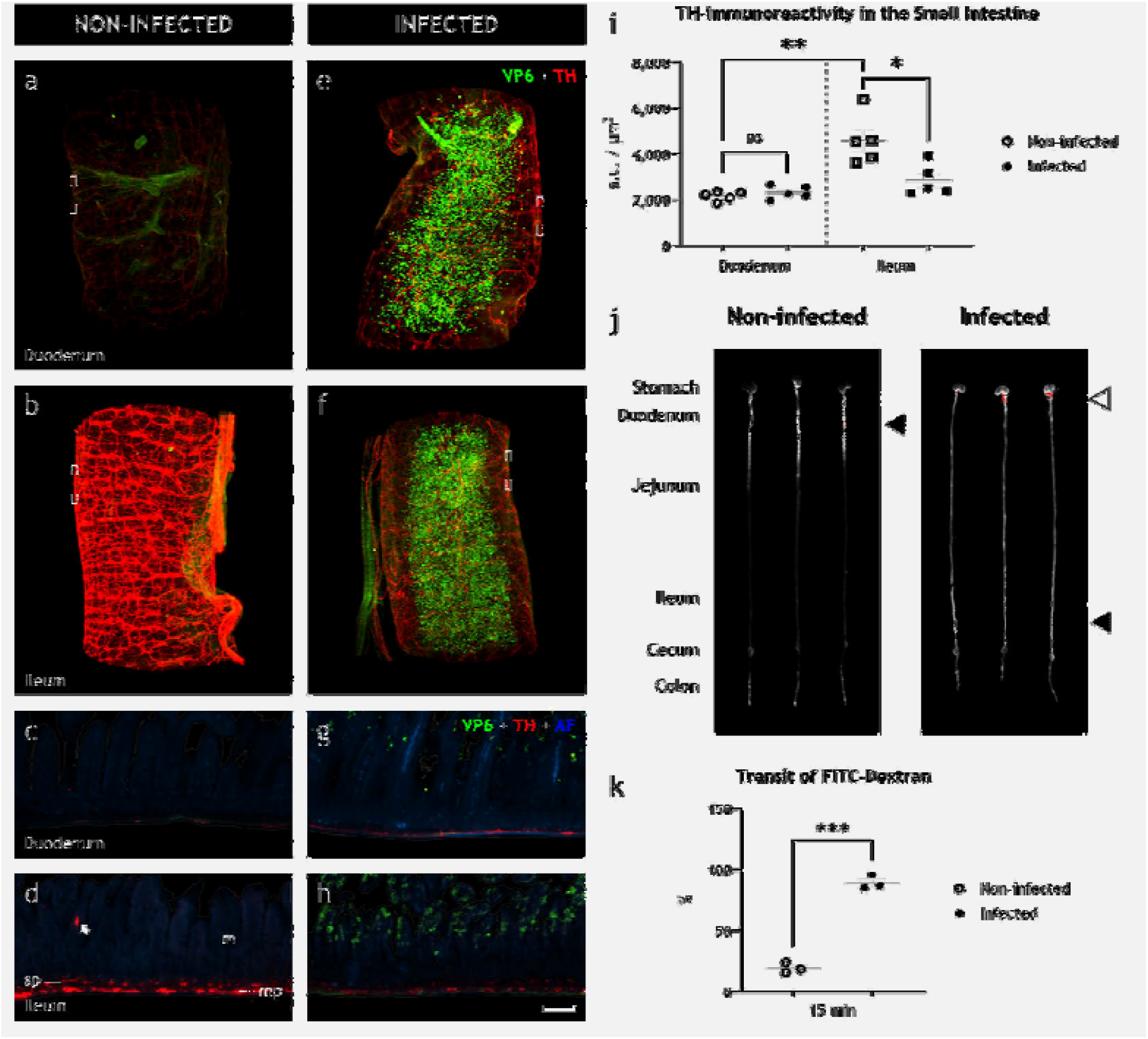
Rotavirus infection simultaneously leads to increased intestinal transit and downregulation of the sympathetic noradrenergic neurons of the autonomic nervous system. Light-sheet micrograph stacks (**a, b, e, f**) and single optical slice (**c, d, g, h**) of infected (right) and non-infected (left) duodenum (**a, c, e, g**) and ileum (**b, d, f, h**) stained for rotavirus VP6 (green) and TH (red), used as a marker for detecting sympathetic axons. Tissue was visualized with autofluorescence (AF; blue). Micrographs in (**c, d, g, h**) correspond to boxed regions in (**a, b, e, f**). An example dopaminergic enteric cell in (**d**), marked with 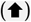, was excluded from the analysis. Note the reduced level of TH immunoreactivity in infected (**f, h**) *vs*. non-infected (**b, d**) ileum. Quantification of TH immunoreactivity (**i**), statistically analysed with two-tailed unpaired (infected *vs*. non-infected) and paired (duodenum *vs*. ileum) *t*-tests, showed no significant difference (ns; p = 0.3236) in the duodenum of infected and non-infected animals, a significant increase in the ileum compared to the duodenum of non-infected animals (**p = 0.0066), and a significant decrease in the ileum of infected compared to non-infected animals (* p = 0.0157). Ultraviolet spectrophotometry of the gastrointestinal tracts of non-infected and 16 h p.i. infant mice 15 min after FITC-dextran treatment (**j**). The average travel distance is marked with (◄); FITC- dextran remnants in the stomach are marked with (<l). Transit of FITC-dextran relative to the entire length of the intestine statistically analysed with the two-tailed unpaired *t*-test with Welch’s correction (**k**) showed a significant (***p = 0.0001) increase in the intestinal motility of the infected animals. m, mucosa; mp, myenteric plexus; sp, submucosal plexus. Scale bar in (**h**) = 100 µm for (**e–h**).

The immunoreactivity in the ileum of the infected animals (n = 5), however, was 2850 ± 309 au/µm^3^. Hence, rotavirus infection led to a significant decrease of TH immunoreactivity in the ileum, but not in the duodenum (see Figure 2a–i). Relative to the average immunoreactivity levels of the uninfected animals, we observed this decrease, which ranged 15–50%, in all infected animals (Figure S2). These data show that rotavirus infection causes robust downregulation of the sympathetic nervous system innervating the ileum.

### Downregulation of the sympathetic nervous system is concomitant with increased intestinal motility

As intestinal motility can be both increased and decreased by the autonomic nervous system (O. Lundgren, 2000; Sharkey & Pittman, 1996; Wood et al., 1999), we next investigated if the rotavirus-induced alteration of the sympathetic nervous system was concomitant with altered intestinal motility *in vivo* by utilizing the well-established fluorescein isothiocyanate (FITC)-dextran intestinal transit model (Hagbom et al., 2020; Istrate et al., 2014). Spectro photographs of resected intestines from animals 16 h post-infection (h p.i.), which had received oral FITC-dextran 15 min prior to termination, clearly showed increased FITC-dextran transit in infected *vs*. non-infected animals (Figure 2j).

The estimated mean relative transit distance (Figure 2k) was 19.1% in the non-infected animals (n = 3) and 89.2% in the infected animals (n = 3). Hence, the infected animals exhibited statistically significantly increased intestinal motility (p = 0.0001) concomitant with reduced sympathetic activity. Notably, the infected animals also showed signs of delayed gastric emptying, visualized by high amounts of remnant FITC-dextran in the stomach (Figure 2j).

### Oral rotavirus infection modulates discrete regions of the brain

The cell bodies of postganglionic sympathetic neurons that innervate the small intestine wall are located in the prevertebral ganglia (Jänig, 1988; Mann & Bell, 1993; Trudrung, Furness, Pompolo, & Messenger, 1994) and receive innervation from the CNS (Berthoud & Powley, 1996; Trudrung et al., 1994). Therefore, we hypothesized that the increased intestinal motility associated with the downregulation of sympathetic nerves during rotavirus infection might be partly controlled by the CNS. To address this question, we investigated the brains of infected and non-infected adult mice using immunohistochemistry for markers of nerve activity.

First, ribosomal protein S6, whose phosphorylated state (pS6) is emblematic of active neurons and parallels expression of the immediate early gene c*Fos* (Knight et al., 2012), was investigated throughout the entire brain. Although a full rostro-caudal survey of the brains of the infected and non-infected animals revealed few differences in the immunoreactivity pattern of pS6 (see *e*.*g*. Figure 3a, b), we observed a significant (p = 0.0233) increase in pS6 immunoreactivity in the area postrema of the infected animals at 48 h p.i. (Figure 3c–f). Within the area postrema, number of pS6-immunoreactive cells per section significantly increased (p = 0.0233) from 21.3 ± 9.2 in non-infected animals (n = 3) to 154.7 ± 47.7 in infected animals (n = 3).

**Figure 3.**
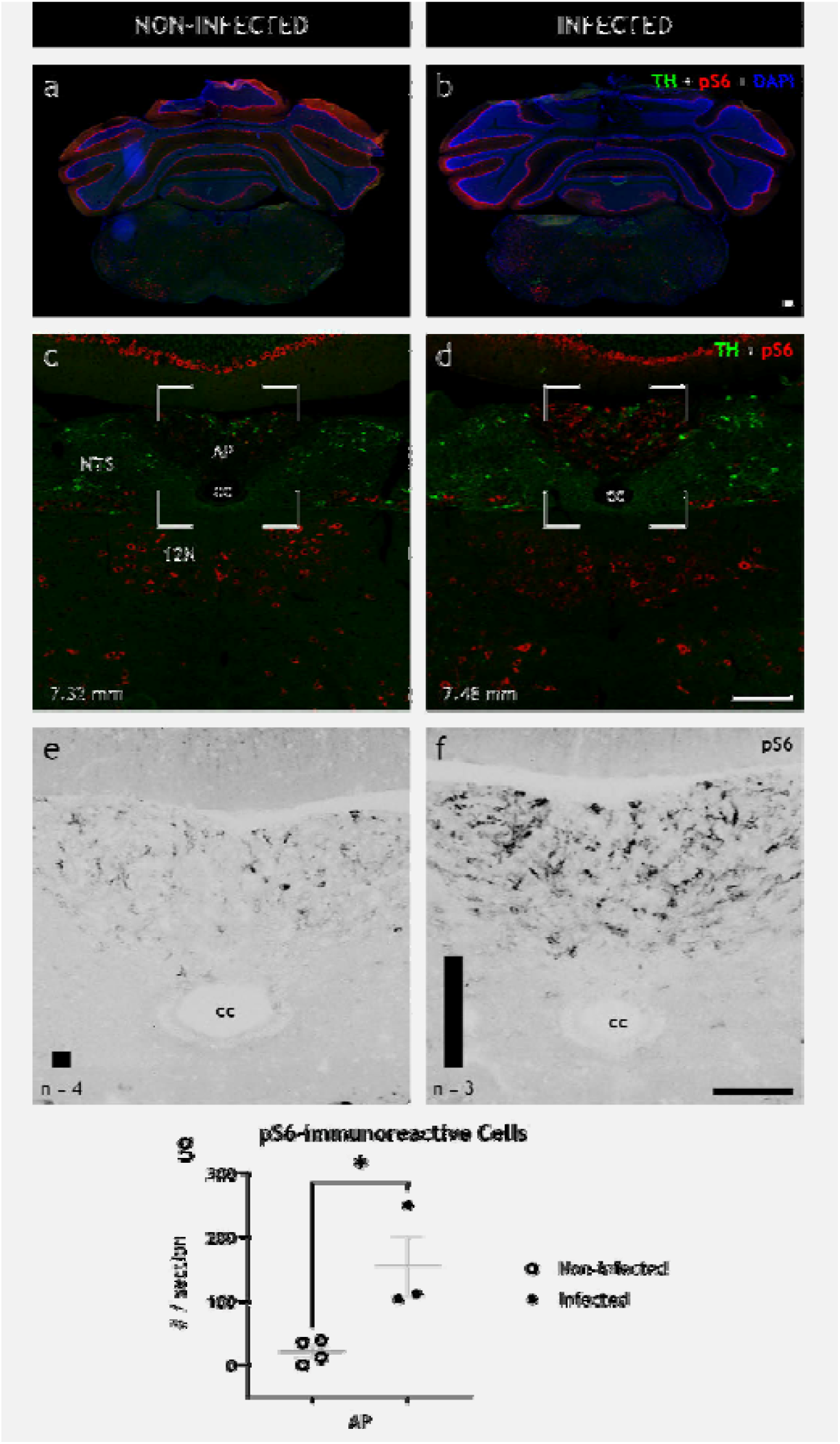
The brainstem is activated by peripheral gastrointestinal rotavirus infection. Representative low- (**a, b**) and high-power (**c, d**) fluorescence micrographs of infected (right) and non-infected (left) coronal brain sections of adult BALB/c female mice immunostained for TH (green), pS6 (red), and DAPI (blue). Magnification of boxed region in (**c, d**) in (**e, f**) only showing pS6; it has been recolored in grayscale for better contrast. Automated quantification of the number of pS6 immunoreactive cells is depicted with a bar in the lower left corner of (**e**) and (**f**) for non- infected and infected animals, respectively. Note the increased level of pS6 immunoreactivity, a marker of activated neurons, in infected animals (**d, f**) but not in non-infected animals (**c, e**). Quantification of pS6 (**g**), statistically analysed with the unpaired t-test, show significant increase (p = 0.0233) of pS6 immunoreactive cell somata in AP. Bregma levels are indicated in the lower left corner. 12N, hypoglossal nucleus; AP, area postrema; cc, central canal; NTS, nucleus of the solitary tract. Scale bar in (**b**) = 100 µm for (**a, b**), in (**d**) = 100 µm for (**c, d**), and in (**f**) = 50 µm for (**e, f**).

As immediate early genes such as *cFos*, and likewise phosphorylation of S6, mark activation in short time frames (hours), while rotavirus infection lasts for days, we next investigated evidence for transcriptional modulations in select brain areas known to control endocrine and autonomic nervous systems. Members of the signal transducer and activator of transcription (STAT) protein family are primarily phosphorylated by the activation of Janus kinase-associated membrane receptors, and the activation of several hypothalamic pathways, particularly regarding feeding behaviour (Furigo, Ramos-Lobo, Frazão, & Donato, 2016), is associated with phosphorylated STAT5 (pSTAT5). We therefore investigated the number of cells expressing pSTAT5 in various brain areas of infected and non-infected animals.

We found pSTAT5 immunoreactive cell somata (Figure 4) in the bed nucleus of stria terminalis (BNST) of all non-infected animals (n = 4) with an average of 6.8 ± 1.7 cells per 14-µm section. Conversely, in the BNST of infected animals (n = 4), we observed a complete and robust absence (p = 0.0286) of pSTAT5 immunoreactive cells (Figure 4). No significant difference was observed in pSTAT5-expressing hypothalamic nuclei, including the arcuate, paraventricular, and periventricular nuclei, as well as the medial preoptic and the anteroventral periventricular areas (Figure S3). Notably, some of these regions showed a high degree of variability among the animals.

**Figure 4.**
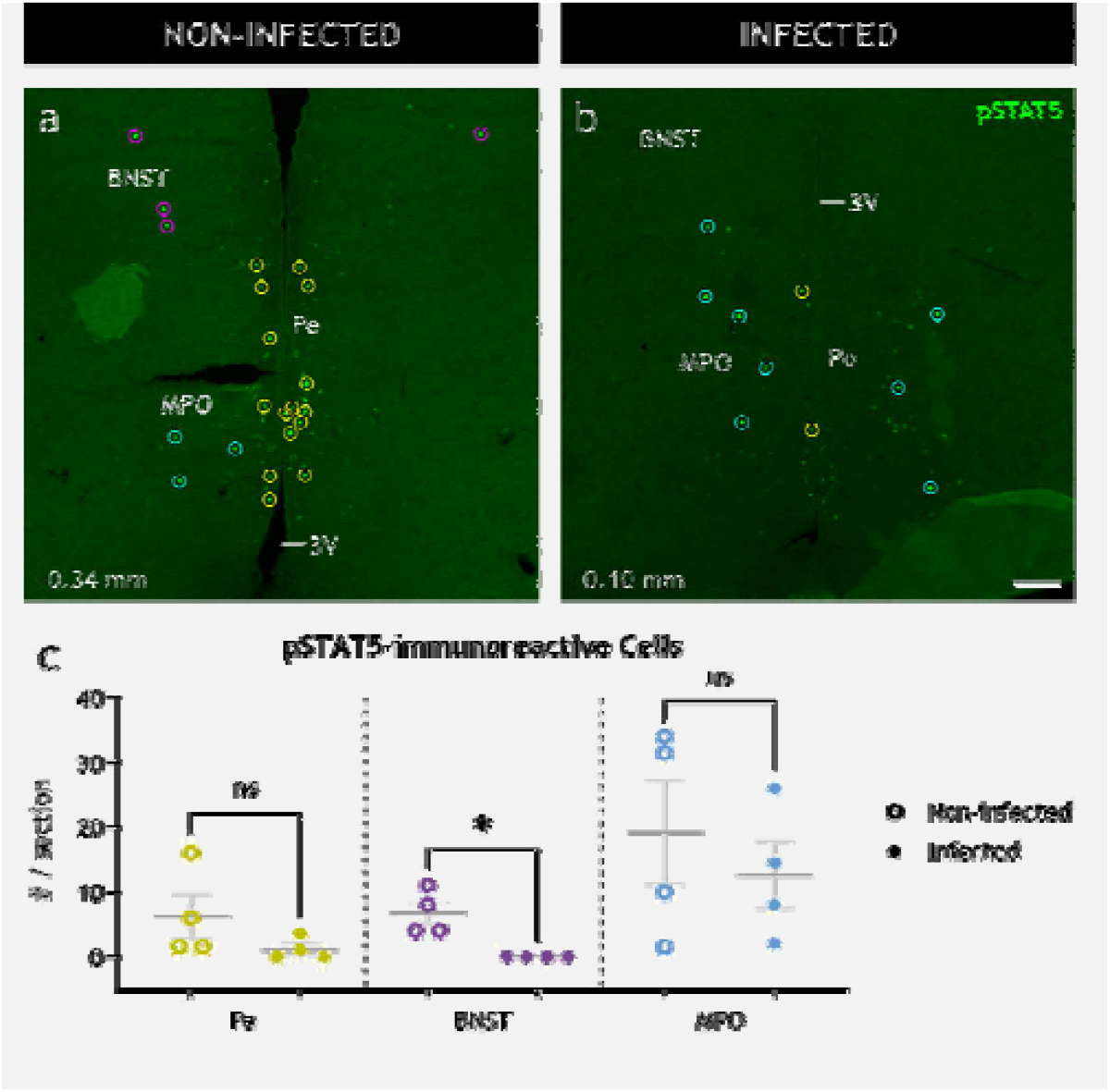
Peripheral gastrointestinal rotavirus infection modulates distinct neuronal populations in the CNS. Representative low-power confocal micrographs of non-infected (**a**) and infected (**b**) coronal brain sections immunostained for pSTAT5 (green), a marker of activated neurons. The immunoreactive cell somata (enclosed in circles) were detected automatically and registered to the corresponding nucleus manually. Quantification of pSTAT5 immunoreactive cell somata (**c**) was statistically analysed with the two-tailed Mann-Whitney test. Note the significant decrease (p = 0.0286) of pSTAT5 immunoreactive cell somata in the BNST, but not the other regions. Bregma levels are indicated in the lower left corner. 3V, third ventricle; BNST, bed nucleus of stria terminalis; MPO, medial preoptic area; Pe, periventricular hypothalamic nucleus. Scale bar in (**b**) = 100 µm for (**a, b**).

### Rotavirus-induced modulation of the CNS is not caused by brain infection

While our data suggest that rotavirus-induced increase of intestinal motility is associated with nervous gut–brain communication, we cannot completely rule out the idea that the virus can reach the brain via the blood and thereby trigger the CNS. Despite little previous evidence for extramucosal spread of EDIM rotavirus (Uhnoo et al., 1990), and the lack of reports of viremia at 16 h p.i., we investigated this possibility with immunohistochemistry. Full rostro-caudal immunohistochemical investigation of fixed neonatal brains at 16 h p.i. (n = 5) and 48 h p.i. (n = 4) did not revealed any evidence of rotavirus VP6 antigen (Figure 5), nor perfused adult brains (n = 5) at 48 h p.i. (data not shown).

**Figure 5.**
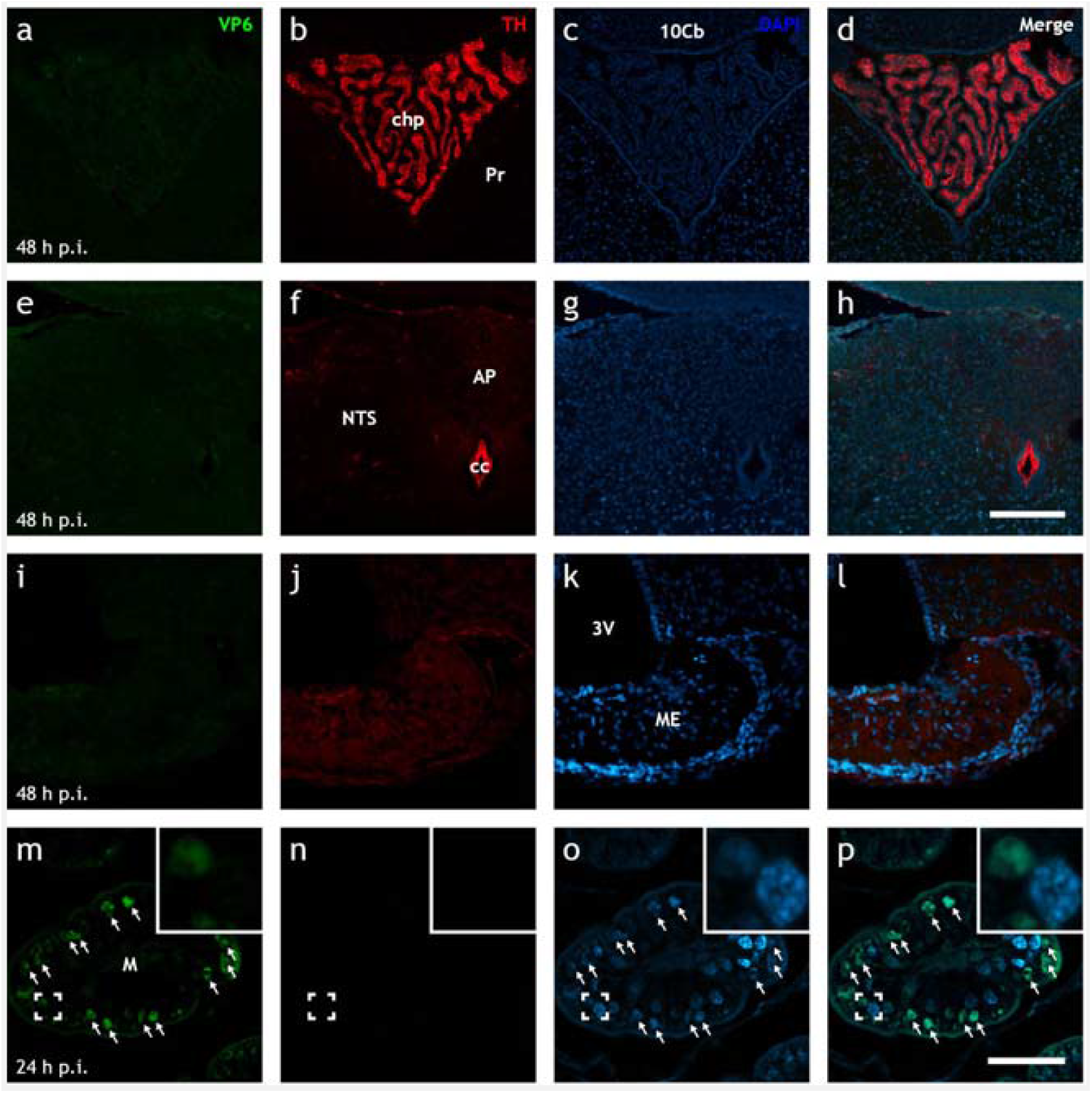
Up to 48 h post infection, EDIM rotavirus is not detected in the brain. Representative low-power Airyscan confocal micrographs of rotavirus-infected neonatal mouse brain (**a–l**) and ileal (**m–p**) sections stained for rotavirus VP6 (green; **a, e, i, m**), TH (red; **b, f, j, n**), and DAPI (blue; **c, g, k, o**). Merge of each row is shown in (**d, h, l, p**). Note the presence of rotavirus-infected cells in the ileum (**↑**; **m–p**) 24 h p.i., but the lack thereof in various areas of the brain as late as 48 h p.i. (**a–l**). 10Cb, 10^th^ lobe of the cerebellum; 3V, third ventricle; AP, area postrema; cc, central canal; chp, choroid plexus; M, mucosa; ME, median eminence; NTS, nucleus of the solitary tract; Pr, prepositus nucleus. Scale bar in (**h**) = 100 µm for (**a–h**) and in (**p**) = 50 µm for (**i–p**).

## Discussion

Previous studies have investigated the mechanisms of rotavirus diarrhoea mainly by focusing on the intrinsic intestinal effects (Ball, Tian, Zeng, Morris, & Estes, 1996; Chang-Graham et al., 2019, 2020; Hagbom et al., 2020, 2011; Istrate et al., 2014; Shirin Kordasti et al., 2006; Ove Lundgren et al., 2000). Although these observations are compelling and have provided important mechanistic information of rotavirus diarrhoea, no information is available on how the gut communicate with CNS before the onset of diarrhoea nor how this communication initiates the illness. By using novel, large-scale volumetric 3D tissue clearing and imaging techniques, we studied the pathophysiology of rotavirus gastroenteritis. We show that rotavirus infection pre-symptomatically disrupts the autonomic balance by downregulating the noradrenergic sympathetic nervous system in ileum, concomitant with increased intestinal transit. In the CNS of infected animals, we found increased pS6 immunoreactivity in the area postrema, and decreased phosphorylated STAT5-immunoreactive neurons in the BNST, which has been associated with autonomic control including stress response. Altogether, these observations reveal that rotavirus signal to CNS before onset of diarrhoea a surprising observation that bring new understanding to how virus give raise to clinical symptoms.

Our 3D illustrations (compare Supplementary Video 1-7) identify a previously unappreciated early widespread infection. Furthermore, our data show that all segments of the small intestine are infected synchronously and demonstrate that the infection triggers neuronal circuitries through the CNS many hours before the development of diarrhoea. These observations are supported clinically, as the well-established early symptoms of rotavirus illness preceding diarrhoea are fever, and nausea/vomiting (stanfordchildrens.org), which are likely to be caused by early gut–brain cross-talks.

The endpoint neurotransmitter of the sympathetic nervous system is noradrenalin (Gershon, 1967; Mann & Bell, 1993). However, as measuring released noradrenalin in the small intestine of infected neonatal mice is challenging due to technical limitations, and released noradrenalin cannot be visualized easily, we chose to investigate the sympathetic system by targeting TH. Since TH is the rate-limiting enzyme of catecholamine biosynthesis (Daubner, Lauriano, Haycock, & Fitzpatrick, 1992; Levitt et al., 1965; Nagatsu et al., 1964), its expression level defines the maximum amount of available neurotransmitter in the cell. Moreover, within the small intestinal wall, TH can only be found in the sympathetic axons (Mann & Bell, 1993), and extrinsic sympathetic denervation of the ileum abolishes all TH immunoreactivity in the intestinal wall. Therefore, our measurements do not appear to be attributed to intrinsic intestinal nerves or any other systems than the sympathetic system. Furthermore, the cell somata of intestinal sympathetic axons receive input from the CNS and are located in the prevertebral ganglia in close proximity to the spinal cord (Jänig, 1988), far from the site of action and shielded from direct viral influence.

By targeting TH, our 3D reconstructions are directly and exclusively correlated to the noradrenergic sympathetic outputs to the small intestine, which we found were downregulated specifically in the ileum, but not in the duodenum, of the rotavirus-infected animals. Occurring within 16 h p.i., this downregulation ranged 15–50% compared to non-infected animals. How this downregulation translates to actual noradrenalin concentration in the cell, and how much noradrenalin is released at the axon terminals, cannot be elucidated from our data. Nonetheless, both clinical data and animal experiments (Istrate et al., 2014) show that the post-infection onset of diarrhoea can vary and occurs between 24 and 48 h.

Notably, we could not find any significant differences in TH immunoreactivity in the duodenum between the infected and non-infected animals, suggesting a tissue-specific rather than general downregulation. Indeed, intestinal segment-specific regulation was reported in 1857 by Eduard Pflüger, who noted that the activation of sympathetic innervation inhibited motility but constricted sphincters (Browning & Travagli, 2014; Jänig, 1988).

The inhibitory effect of the noradrenalin from the sympathetic nervous system on the small intestine is well established (Gershon, 1967; Kadowaki, Yoneda, & Takaki, 2003). Early histochemical investigations have determined that axons of the sympathetic postganglionic neurons are present in the submucosal and myenteric plexuses, and also extend to the villi in the mucosa (Schultzberg et al., 1980). Furthermore, functional and pharmacological studies show that noradrenalin mainly acts on α_1_-adrenergic receptors to excite myenteric neurons and thereby increase intestinal motility (Furuichi et al., 2001; Schemann, 1991). Further, enteric glia both regulate gastrointestinal motility (Gulbransen & Sharkey, 2012) and express adrenergic receptors (Nasser, Ho, & Sharkey, 2006). Indeed, rotavirus activates enteric glia cells via serotonin (Hagbom et al., 2020). Altogether, this suggests that noradrenalin simultaneously acts on enteric neurons and glia cells, parasympathetic axons and smooth muscle cells to coordinately inhibit intestinal motility. In accordance with our data, reducing the available noradrenalin will remove these inhibitions, *i*.*e*. remove the brake, and shift the balance towards increased intestinal motility, as illustrated in Figure 6.

**Figure 6.**
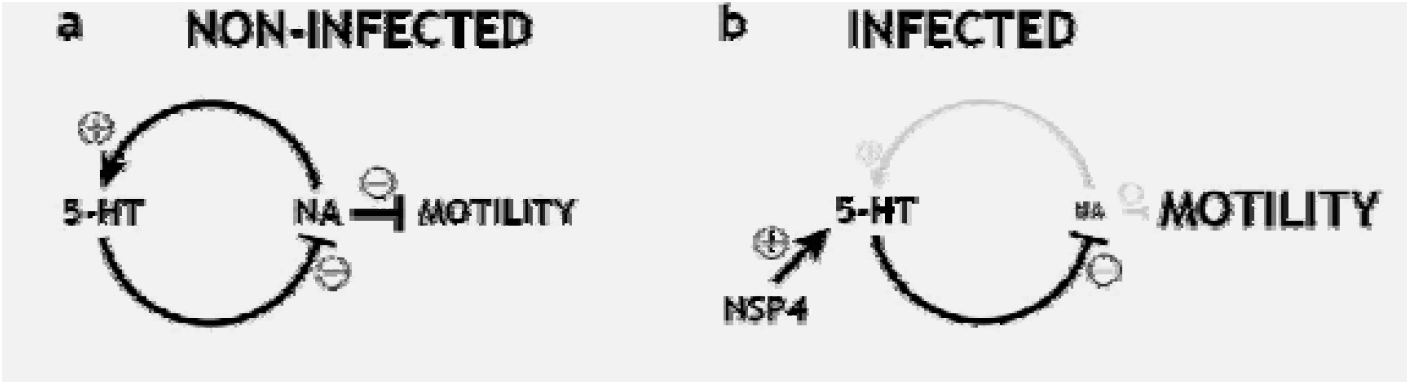
Sympathetic feedback loop is disrupted in ileum during infection. Schematic representation of the sympathetic feedback loop at enterochromaffin (EC) cells. In normal conditions (**a**), sympathetic noradrenalin (NA) will stimulate the release of serotonin (5-HT) from the EC cells, which will act on the ascending vagal pathways to inhibit the excessive release of noradrenalin. During infection, NSP4 will disrupt the autonomic balance by circumventing noradrenalin and inducing the continuous release of serotonin. This will cause reduced sympathetic tone, which will leave the parasympathetic nervous system to stimulate intestinal motility unhindered and cause diarrhoea.

We observed increased pS6 immunoreactivity in area postrema and decreased number of pSTAT5 immunoreactive cell somata in the BNST and no rotavirus antigen in CNS. Based on these observations we conclude that the sympathetic downregulation in the intestine resulted from gut–brain nervous signalling rather than direct infection and/or cytokine stimulation. This conclusion is supported by the fact that rotavirus is associated with limited inflammatory response in both human and mice (Greenberg & Estes, 2009; Hagbom et al., 2021; O. Lundgren, 2000; Ove Lundgren & Svensson, 2001; Morris & Estes, 2001), and that the EDIM murine rotavirus strain used in the present study has not been associated with extramucosal spread earlier than 72 h p.i. (Kraft, 1958) nor hepatic infiltration (Uhnoo et al., 1990).

Abnormal gastric motor function, as manifested by delayed emptying, has been reported in rotavirus-infected children (Bardhan, Salam, & Molla, 1992), and has been proposed to be associated with gastrointestinal hormones, neuronal pathways (including non-cholinergic and non-adrenergic), vagal neurons, and CNS control. The precise mechanisms, remain unresolved (Crawford et al., 2017). The FITC-dextran remnants in the stomach of the infected animals (Figure 2j, k) suggest the occurrence of delayed gastric emptying during the early stages of infection. Together with our other data showing downregulation of the sympathetic nervous system in the ileum, it strengthens the view of nerves participating in rotavirus illnesses. Altogether, our data suggest altered autonomic control as the underlying cause of other symptoms as well, and further investigation of the stomach, for example, is warranted.

Interestingly, we found a strong reduction of pSTAT5 immunoreactive cell somata in the BNST of infected animals (Figure 4). Spinal neuron projections directly to the BNST have been reported (Menétrey & de Pommery, 1991). Further, BNST sends projections to the dorsal motor nucleus of the vagus (Hopkins & Holstege, 1978), the nucleus ambiguous (Holstege, Meiners, & Tan, 1985), and the nucleus of the solitary tract (Hopkins & Holstege, 1978), *i*.*e*., the brain centra involved in controlling gastrointestinal motility (Browning & Travagli, 2014; Gillis, Quest, Pagani, & Norman, 2011). Furthermore, the BNST is involved in several autonomic regulations responding to non-fear-associated stress, and alters both blood pressure (Koikegami, Kimoto, & Kido, 1953) and heart rate. Our data showing modulation of the BNST in response to rotavirus infection strengthens the view of BNST involvement in intestinal motility and possibly symptoms of illness.

EC cells of the small intestine modulate neuronal signalling, including intestinal motility and secretion. Rotavirus, as well as NSP4, stimulates serotonin release from EC cells (Chang-Graham et al., 2019, 2020; Hagbom et al., 2011) and directly modulates ascending vagal pathways (Crawford et al., 2017; Hagbom et al., 2011). EC cells also receive direct sympathetic input, and noradrenalin excites EC cells to release serotonin (Bellono et al., 2017). Based on these reports and our collective observations, we propose EC cells as an intestinal sensor using vagal outputs and sympathetic intestinal sensory feedback to modulate gastrointestinal motility. This proposal provides both molecular and systemic explanations for how rotavirus infection can disrupt the autonomic balance. Furthermore, we suggest the nucleus of the solitary tract, the area postrema, and the BNST as central relay points of this feedback loop (Figure 7).

**Figure 7.**
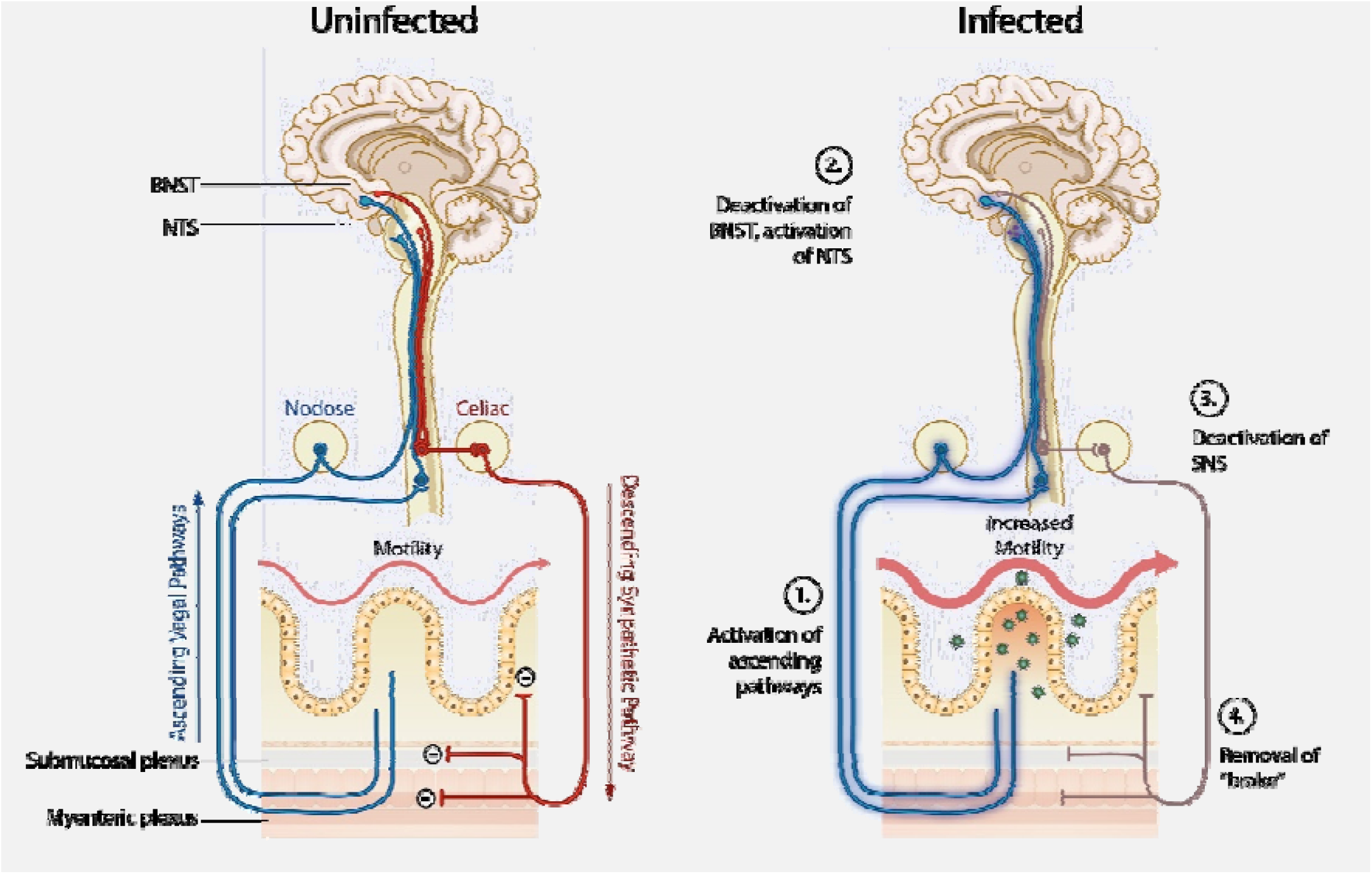
Proposed mechanistic model for rotavirus induced diarrhoea involving gut-brain cross-talk. In normal conditions (**a**), sensory information is relayed to the CNS, which keeps the autonomic nervous system in balance. During infection, rotavirus will cause: (1) excessive release of serotonin and thereby activation of the afferent vagal pathways, which are (2) relayed to the CNS to modulate discrete regions including the BNST and NTS. In the CNS, the signal is processed, and (3) the efferent sympathetic nerves innervating the ileum are downregulated. In the ileum, (4) reduced levels of noradrenalin from the sympathetic nervous system will lead to less inhibition (i.e., removal of the brake) of the enteric neurons, parasympathetic axons, smooth muscle cells, and enteric glia cells, which together shift the autonomic balance towards increased intestinal motility.

## Conclusions

We showed and quantified the extent of rotavirus infection of the small intestine in 3D and identified centrally relayed downregulation of the sympathetic innervation of ileum, concomitant with increased intestinal transit and altered brain activity before onset of diarrhoea. We found increased pS6 immunoreactivity in area postrema and decreased phosphorylated STAT5-immunoreactive neurons in the BNST, which has been associated with autonomic control including stress response. Collectively, our data provide novel information how rotavirus causes illness and communicate with nerves and the brain.

## Acknowledgements

Financial support for this study was provided by the Swedish Research Council (Grants 2014-02827, 2017-01479, 2018-02862).

## Author contributions

A.H., L.S., and M.H. designed the studies; A.H., L.S., and M.H. conducted the experiments; A.H. and M.H. analysed the data; A.H. wrote the manuscript with input from all authors. All authors read and approved the final manuscript.

## Supplementary Videos

**Video 1.**
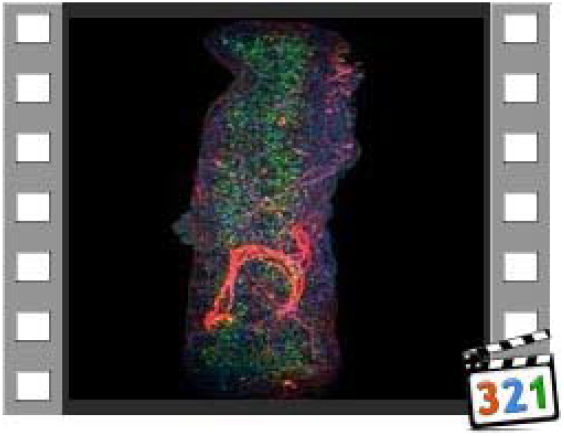
Supplementary video to Figure 1a.

**Video 2.**
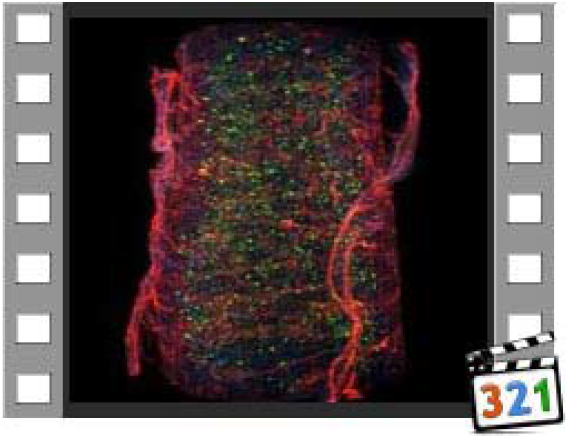
Supplementary video to Figure 1d.

**Video 3.**
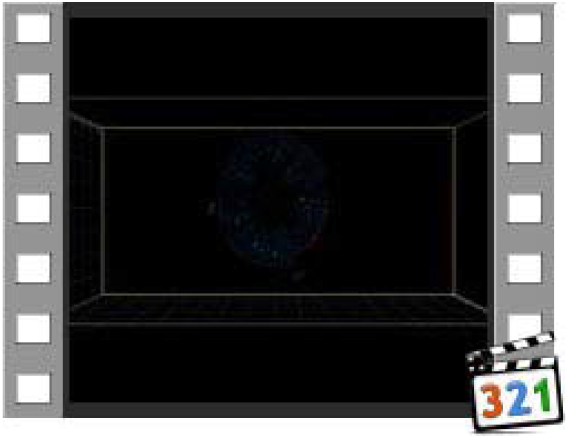
Supplementary video to Figure 1d.

**Video 4.**
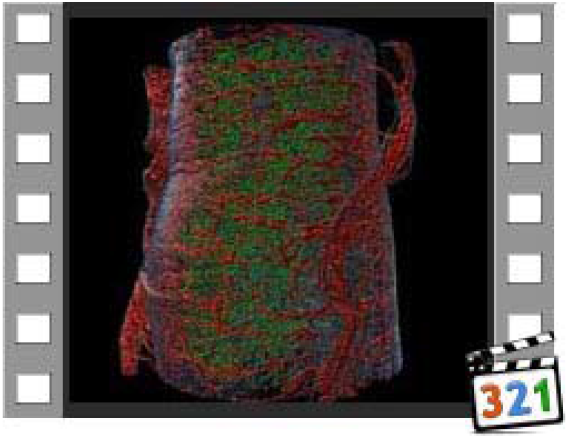
Supplementary video to Figure 1e.

**Video 5.**
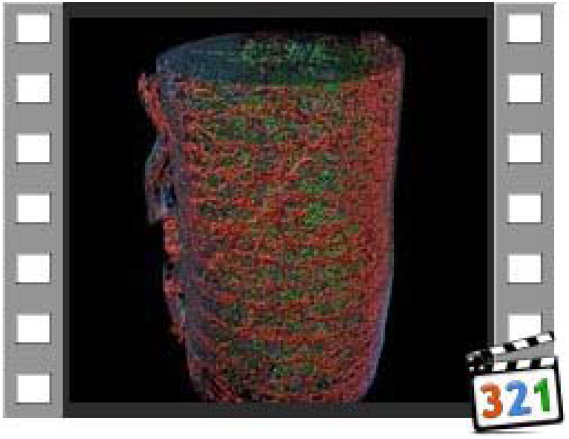
Supplementary video to Figure 1e.

**Video 6.**
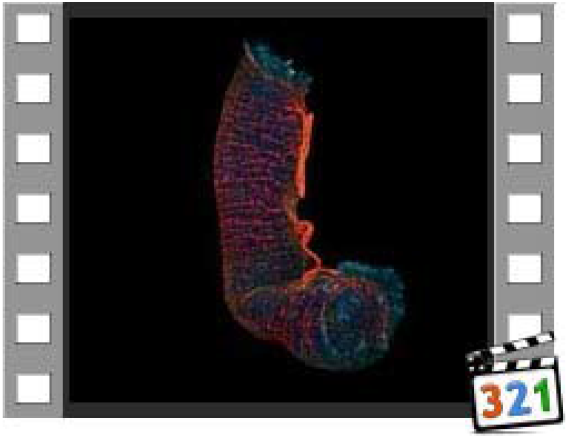
Supplementary video to Figure 2b.

**Video 7.**
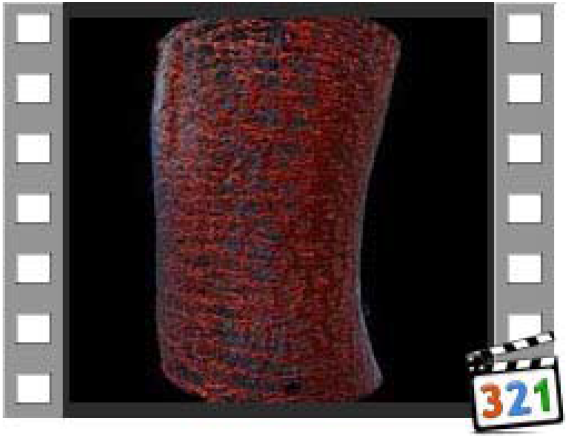
Supplementary video to Figure 2b.

**Figure S1.**
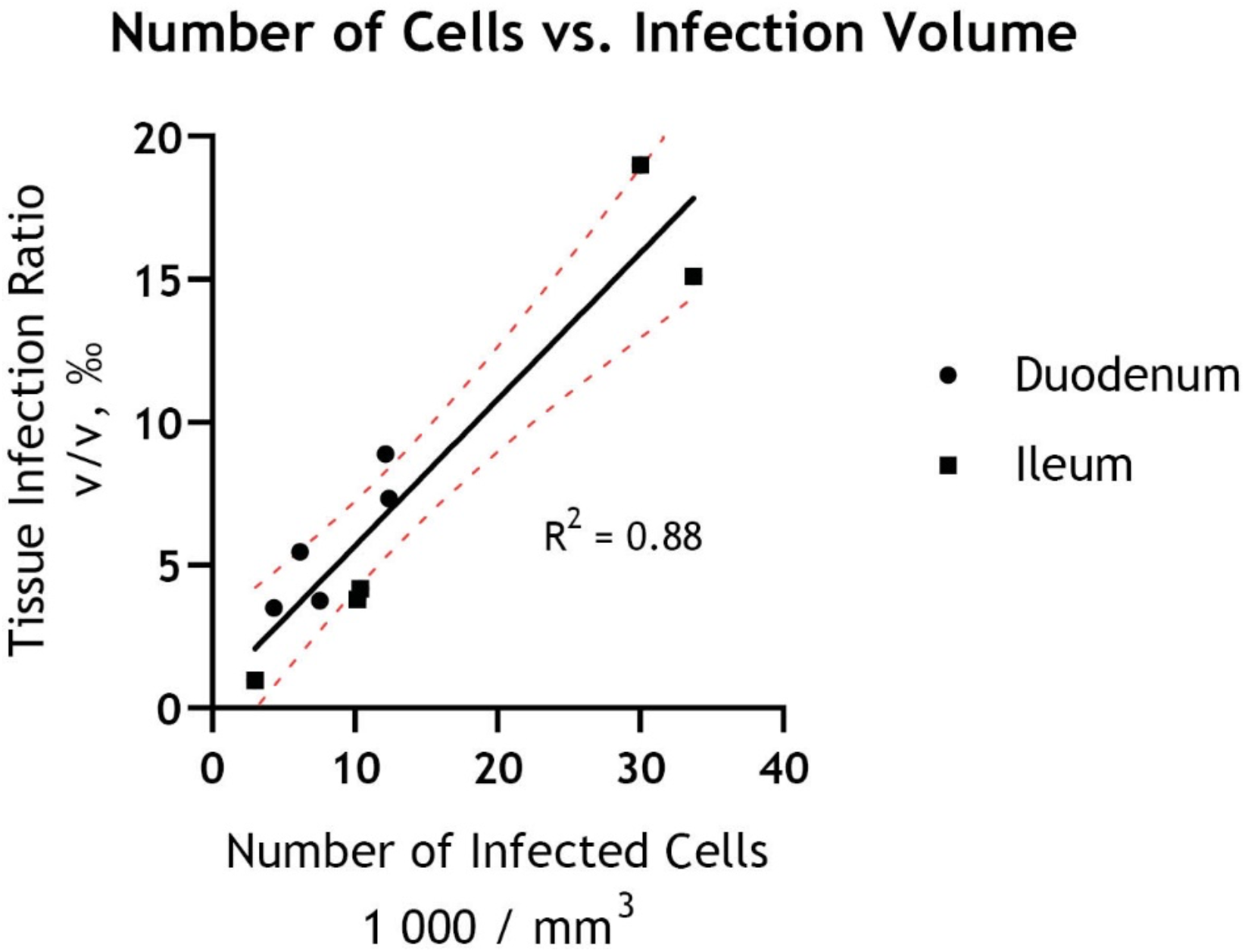
Two different approaches to estimate the degree of infection yields similar results. Estimated relative number of infected cells plotted against tissue infection ratio show a linear relationship between the two approaches.

**Figure S2.**
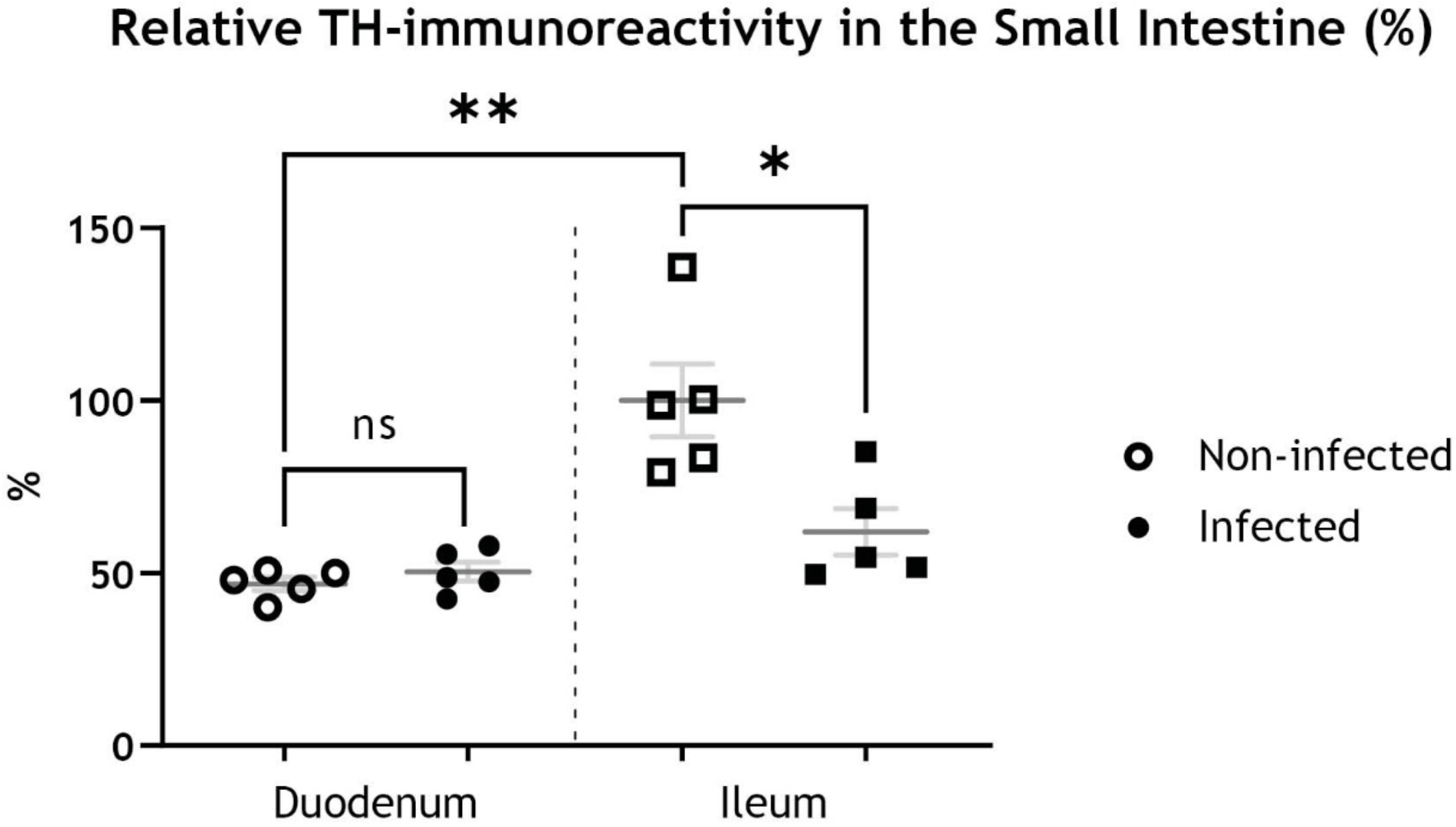
Rotavirus infection reduces the amount of ileal TH-immunoreactivity. Supplementary graph to Figure 2. Quantification of relative TH-immunoreactivity, statistically analysed with two-tailed unpaired (infected vs. non-infected) and paired (duodenum vs. ileum) t-tests show no significant (ns; p = 0.3236) difference in duodenum of infected and non-infected animals, significant increase in ileum compared to duodenum of non-infected animals (**; p = 0.0066) and significant decrease in ileum of infected compared to non-infected animals (*; p = 0.0157). Data presented relative to average TH-immunoreactivity in non-infected ileum.

**Figure S3.**
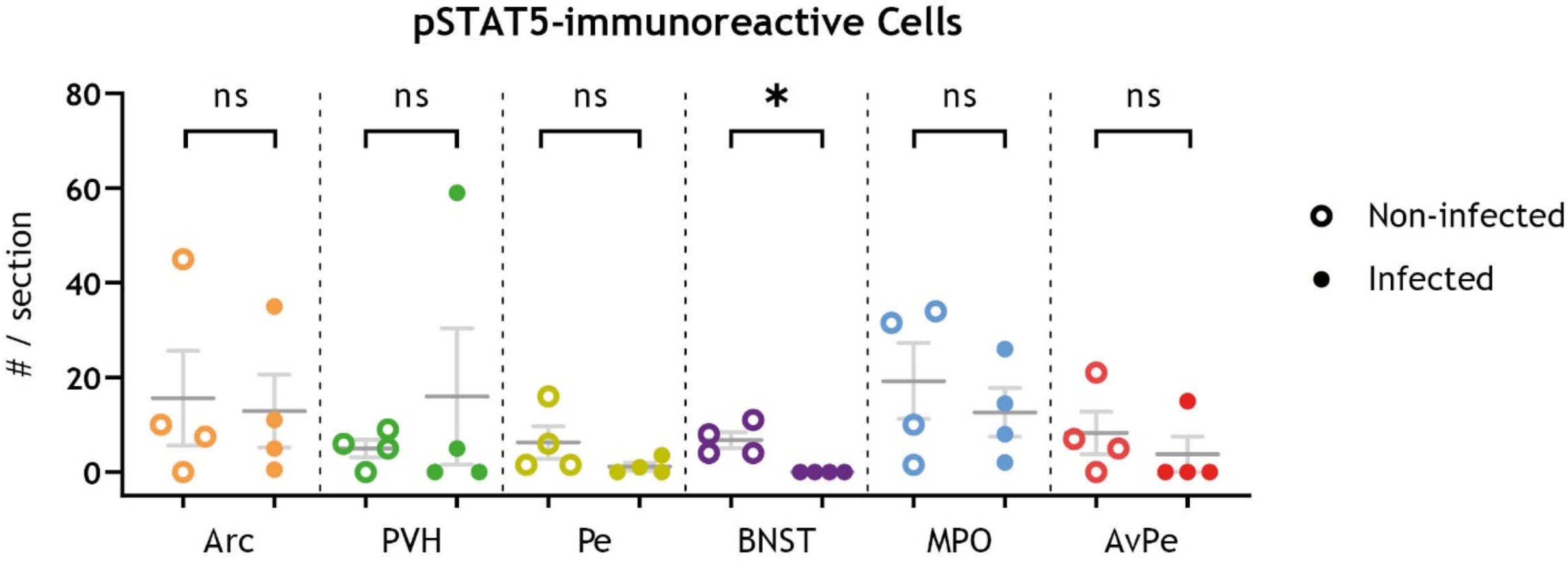
Peripheral gastrointestinal rotavirus infection modulates pSTAT5 in the BNST but no hypothalamic regions. Supplementary data to Figure 4. Quantification of pSTAT5-immunoreactive cell somata from infected and non-infected animals, statistically analysed with two-tailed Mann-Whitney test, identifies a decrease in the BNST (p = 0.0286) but no significant difference in various hypothalamic regions. Arc, arcuate nucleus of hypothalamus; AvPe, anteroventral periventricular nucleus; BNST, bed nucleus of stria terminalis; MPO, medial preoptic area; NTS, nucleus of the solitary tract; Pe, periventricular hypothalamic nucleus; pSTAT5, phosphorylated transducer and activator of transcription 5; PVH, paraventricular nucleus of hypothalamus.

**Figure S4.**
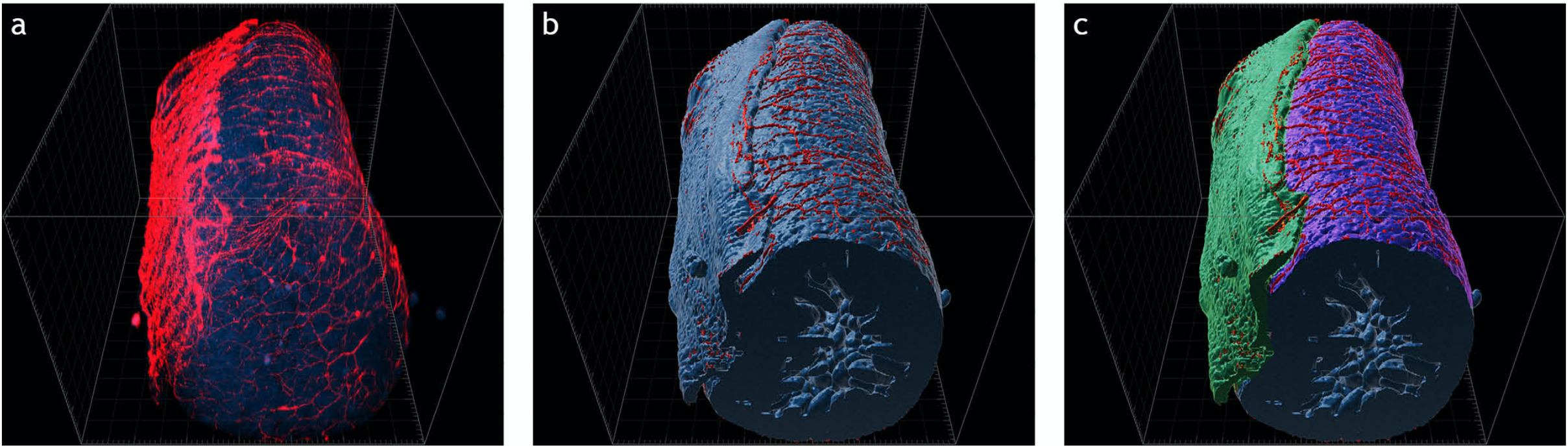
3D investigation reveals damaged tissues that can be excluded from analysis. Maximum intensity projection of light sheet micrograph stacks (a) from mouse ileum stained for tyrosine hydroxylase (TH; red) to mark sympathetic innervation of the intestine. Tissue visualized with autofluorescence (AF; blue). 3D surface reconstruction from (a) in (b, c). Muscularis (green) and submucosa (purple) pseudo-colored in (c). Note damage to the outer layer of the intestinal wall leaving the submucosal plexus exposed. Lack of myenteric plexus from tissue sample is hidden to the naked eye, barely detectable on micrographs, but fully identified with 3D surface modelling.

